# Association of poor virus specific immunoglobulin G antibody responses with higher viral load is seen in Bangladeshi pregnant women having acute Hepatitis E Genotype 1 infection

**DOI:** 10.1101/2020.06.12.148528

**Authors:** Rosy Sultana, Md Tarikul Islam, Golam Sarower Bhuyan, Farjana Akther Noor, Suprovath Kumar Sarker, Noorjahan Maliha, Zahid Hassan, Mohabbat Hossain, Mamunur Rashid, K Zaman, Muhammad Manjurul Karim, Firdausi Qadri, Kaiissar Mannoor

**Author notes:** **Corresponding author** (KM). **Disclosures** All the authors declare that they have no competing interests.

## Abstract

Although Hepatitis E viral illness is usually self-limiting, higher rates of morbidity and mortality are frequently observed during pregnancy in South Asian countries including Bangladesh. Of the four common variants, hepatitis E virus genotype 1 is mainly prevalent in South Asian countries. Pregnant women usually suffer from a state of immunosuppression. It is yet to be known whether virus specific immunoglobulin G (IgG) immune responses have any association with the vulnerability of pregnant women to acute hepatitis with E virus. The study aimed to compare the viral load and IgG responses of hepatitis E-infected pregnant women with that of non-pregnant women with same infection. Real Time –quantitative reverse transcription Polymerase Chain Reaction and Sanger sequencing were performed to determine the viral load and genotype, respectively, whereas Enzyme Linked Immunosorbent Assay method was used to determine hepatitis E virus specific serum IgG antibody index along with IgG avidity index. Although significant negative correlations were observed between log viral copy number and log IgG antibody index in the late acute phases of jaundice for both pregnant (r= −0.7971, p=0.0002) and non-pregnant women (r= −0.9117, p=0.0002), serum log viral copy number of pregnant women was significantly higher than that of the non-pregnant counterpart (p=0.0196) in the late acute stage of jaundice. In addition, log hepatitis E virus IgG antibody index of pregnant women was significantly lower than the non-pregnant women in the late phase of jaundice induced by hepatitis E virus (p=0.0303). Moreover, pregnant women with acute hepatitis E had higher cross-reactive IgG than in the non-pregnant women (p=0.0017). All the patients got infected with hepatitis E virus were in Genotype 1 variety. The study demonstrates that virus-specific poor IgG responses might be responsible for vulnerability of pregnant women to acute hepatitis with hepatitis E virus.

**Author Summary:** Acute hepatitis caused by hepatitis E virus (HEV) Genotype 1 is a public health problem in Asian countries and especially it poses a potential health threat to pregnant women causing 19% to 25% mortality, particularly in South Asian countries including Bangladesh. The study aimed to explore whether HEV IgG immune responses were compromised during pregnancy, which might contribute to higher viral load and disease severity. Accordingly, pregnant and non-pregnant women with acute hepatitis (clinically presented with nausea, loss of appetite and /or jaundice) were enrolled from different tertiary care hospitals in Dhaka city. All these patients were screened and hepatitis E were differentiated from other hepatitis (caused by A, B, C) using Enzyme Linked Immunosorbent Assay (ELISA) methods. HEV IgG antibody/avidity indices and viral loads were measured using ELISA and real time quantitative polymerase chain reaction (RT-qPCR), respectively. The study showed that pregnant women with acute hepatitis E had lower IgG indices with higher viral load than their non-pregnant counterpart. Overall, the study revealed that virus-specific poor IgG responses might render pregnant women vulnerable to acute hepatitis E of varying degree of severity which might be associated with higher viral load.

## Introduction

Acute hepatitis caused by HEV is a global public health concern as it affects 20 million individuals with an annual mortality of 44,000 worldwide [1]. The disease is usually self-limiting in men and non-pregnant women with a case-fatality rate of <0.1% [2]. However, severity is documented among pregnant women in Bangladesh, often leading to deaths in up to 19–25% of the cases [3]. During the last decades, an improved understanding of the natural history of HEV infection, reservoirs, and transmissions modes has been achieved [4]. Moreover, studies have reported significant changes in immunological and hormonal responses in pregnant women with fulminant hepatic failure caused by HEV, explaining high morbidity and mortality rate in this group of patients [2]. However, it is not well understood why there are disparities in terms of disease severity and outcomes in some geographical locations in developing countries and in pregnant women with HEV Genotype 1 hepatitis. There are some reports demonstrating high viral load in pregnant women with acute hepatitis compared to that of non-pregnant [5, 6].

The determination of anti-HEV specific antibodies (IgM and IgG) and/or detection of viral RNA in serum are the main diagnostic methods for HEV diagnosis. HEV-RNA appears in the blood shortly before the onset of symptoms and generally peaks during early acute phase of illness with persistent viremia for approximately 3-6 weeks [4, 7]. While anti-HEV IgM response is an indicator of acute infection, peak of anti-HEV IgG predicts convalescent phase ensuring recovery of the patients [8], although acute and ongoing infection is often accompanied by both anti-HEV IgM and IgG [9]. Currently, no kits are commercially available for quantitative measurement of anti-HEV IgG antibody level. However, HEV IgG SI (sample index) value, antibody avidity index and their correlation with the viral load can be used as an important tool for prediction of viral elimination and assessment of immune response against the virus [10]. Moreover, studies on immune responses among individuals with HEV hepatitis as well as various animal models on HEV infections reported a protective role of IgG antibody [11]. In humans, HEV infections had been reported to promote Th1 immunity, whereas Th2 immune responses are induced during the pregnancy to ensure the safety of the fetus [2]. An imbalance between Th1 and Th2 responses was expected when pregnant women are infected with HEV, which may suppress anti-HEV IgG immune responses in pregnancy. This study was carried out to investigate whether there were any differences in IgG immune responses imparted by anti-HEV IgG between pregnant and non-pregnant Bangladeshi women with icteric symptoms due to acute HEV hepatitis.

## Materials and Methods

### Study design, patient selection and data collection

This cross-sectional study spanned between August 2016 to September 2019 which recruited a total 153 (67 pregnant, 86 non-pregnant) consecutively admitted suspected acute hepatitis female patients, age range 18-50 years, from Female Medicine and Antenatal wards of three tertiary care hospitals of Dhaka, namely Dhaka Medical College Hospital, Bangabandhu Sheikh Mujib Medical University Hospital and Bangladesh Institute of Health Sciences and General Hospital. Recruitment criteria included, diagnosis of acute viral hepatitis was confirmed by the presence of constellation of clinical symptoms (nausea, anorexia, upper abdominal pain, low grade fever and yellowish urine), presence of jaundice for <6 weeks, and biochemical evidence of hepatocellular damage demonstrated by high levels of alanine transaminase (ALT) and aspartate transaminase (AST) or both. An informed written consent was taken from each patient. A structured questionnaire was filled out which included information on each patient’s socio-demographic particulars (age, residence, annual family income, education level etc.), potential risk factors for viral transmission (source of drinking water, living arrangements, food habits etc.), and previous history of jaundice. The study was approved by the Institutional Research Review Boards of Bangladesh University of Health Sciences (BUHS) and Dhaka Medical College (DMC), Dhaka, Bangladesh.

### Specimen collection and serum preparation

For the study, 4.0 ml blood sample was collected in a plain vacutainer tube from each participant and transported to the institute for developing Science and Health initiatives (ideSHi) laboratory in a cool (4-8°C) box within an earliest possible convenience. Serum was prepared after centrifugation of the sample tube for 10 minutes at 3000 r.p.m. in a refrigerated Centrifuge machine. Serum was aliquoted (800 microliter in each of the three microcentrifuge tubes). One aliquot was used for biochemical liver function tests, another for serological screening of viral hepatitis markers on the same day of collection and third aliquot was preserved at −80 °C for molecular analysis at later period.

### Liver Function Tests, anti-HEV IgM and anti-HEV IgG Detection

Liver function tests; serum bilirubin, ALT and AST were performed using a biochemistry autoanalyzer (Dimension, Siemens, Germany). Patients with serum bilirubin level ≥1.2 mg/dl of blood and ALT level ≥ 40 IU/L were subjected to screening for viral hepatitis markers, such as Hepatitis B surface antigens (HBsAg) and Hepatitis C antibodies (anti-HCV) using ELISA kits (BioKits, Spain), IgM antibodies for Hepatitis A (anti-HAV IgM) and anti-HEV IgM and IgG using ELISA kits (Wantai, China). To determine anti-HEV IgG index, a minor modification of the manufacturer’s protocol was adopted. The IgG sample index (SI) value was calculated as follows: [(Sample absorbance-Negative Control absorbance)/(Positive control absorbance-Negative control absorbance)] *dilution factor *100.

### Avidity ELISA

Avidity ELISA was performed using the commercially available anti-HEV IgG antibody (Wantai, China) kit with minor modifications. The samples were incubated before addition of conjugate with 7M urea solution in some wells of the HEV antigen coated plates, while the same samples in another set of wells were treated with buffer following the protocol provided with the kit. Finally, the avidity index was calculated using the following formula: Avidity Index (AI) = (Sample OD of untreated well-sample OD of urea treated well)*100/ Sample OD of untreated well.

### Categorization of the Study participants

The hepatitis A, B and C positive samples were excluded and finally the study analyzed 121 (57 pregnant, 64 non-pregnant) patients with acute HEV hepatitis. On the basis of pregnancy status and duration of jaundice, the patients were divided into (1) early acute phase, ≤14 days (2) late acute phase≥14 days) [12] of jaundice.

### Determination of Viral Genotype and Copy Number

To calculate the viral load, seven synthetic viral nucleic acid standards containing the conserved region of HEV genome of four common genotypes with known copy number were used. A total of 78 serum samples (positive for HEV IgM in ELISA) were subjected to RNA extraction using QIAamp Viral RNA Mini Kit (Qiagen, Germany). Using SuperScript™ III One-Step RT-PCR System with Platinum™ Taq (Invitrogen, USA) and specific primers targeting the conserved regions of the ORF2 of all HEV genotypes, real time RT-PCR was run on CFX96 Touch™ Real-Time PCR detection system (Bio-Rad). Viral load was calculated using the cycle of threshold (**Ct)** values. For genotype determination of the purified PCR products, Sanger nucleotide sequencing was carried out using the ABI PRISM 310 Automated Sequencer (Applied Biosystems, USA) and the sequences were analyzed using BLAST tool to compare the query sequence with the reference sequence retrieved from NCBI database.

### Statistical Analysis

The IgG sample index, standard curve for viral copy number and avidity index were calculated using Microsoft Excel v2016. Chi-square test with 95% confidence interval (CI) using SPSS v17.0 was performed to find the association among different socio-demographic parameters. Pearson correlation analysis was done for HEV copy number and HEV IgG SI values of pregnant and non-pregnant women distributed in early and late acute phases of jaundice. Age, duration of jaundice, serum bilirubin, ALT, AST, viral load and IgG SI values were compared between different group stratifications using unpaired Student’s t text and Mann–Whitney test, as appropriate. The analyses were performed using GraphPad Prism v7.0 and a p-value of less than 0.05 was considered statistically significant.

## Results

### Sociodemographic and biochemical profiles

A total of 121 women with acute HEV infection (57 pregnant and 64 non pregnant) were included for data generation and analysis. Of the 121 patients, 78 were positive for anti-HEV IgM antibody, which is an early marker of acute hepatitis E, whereas the other 43 were negative for anti-HEV IgM but positive for HEV IgG antibody. Distribution of these patients on the basis of residence, housing type, habit of eating street food, education and per-capita income did not show statistically significant association with HEV IgM positivity (p=0.302, 0.186, 0.434, 0.093 and 0.474, respectively). Conversely, shared facilities for living (p=0.006, OR= 4.47, 95% CI: 1.53 – 13.06) and unsafe drinking water (p=0.017, OR= 3.26, 95% CI: 1.23 – 8.62) (Table 1) showed statistically significant association with HEV IgM positivity.

**Table 1.** Socio-demographic status of the HEV IgM positive and negative hepatitis patients (N=121)

| Variables | HEVIgM+ve<br>(n=78) | HEVIgM-ve<br>(n=43) | P value | Odds<br>Ratio | 95%<br>Lower | CI<br>Upper |
| --- | --- | --- | --- | --- | --- | --- |
| **Residence** |  |  |  |  |  |  |
| Semi-urban (29) | 16 (55.2) | 13 (44.8) | 0.302 | 1.67 | 0.63 | 4.41 |
| Urban (92) | 62 (67.4) | 30 (32.6) |  |  |  |  |
| **Housing** |  |  |  |  |  |  |
| Tin shade (62) | 39 (62.9) | 23 (37.1) | 0.186 | 1.98 | 0.72 | 5.48 |
| Building (59) | 39 (66.1) | 20 (33.9) |  |  |  |  |
| **Living arrangements** |  |  |  |  |  |  |
| Individual facilities (86) | 49 (57.0) | 37 (43.0) | 0.006 | 4.47 | 1.53 | 13.06 |
| Shared facilities (35) | 29 (82.9) | 6 (17.1) |  |  |  |  |
| **Drinking water** |  |  |  |  |  |  |
| *Safe drinking (38) | 19 (50.0) | 19 (50.0) | 0.017 | 3.26 | 1.23 | 8.62 |
| **Unsafe drinking (83) | 59 (71.1) | 24 (28.9) |  |  |  |  |
| **Street food habit** |  |  |  |  |  |  |
| Yes (102) | 65 (63.7) | 37 (36.3) | 0.434 | 0.63 | 0.19 | 2.03 |
| No (19) | 13 (68.4) | 6 (31.6) |  |  |  |  |
| **Educational status** |  |  |  |  |  |  |
| Up to twelve grade (102) | 66 (64.7) | 36 (35.3) | 0.093 | 2.18 | 0.88 | 5.39 |
| Graduate and higher (19) | 12 (63.2) | 7 (36.8) |  |  |  |  |
| **Per-capita income (USD)<sup>§</sup>** |  |  |  |  |  |  |
| Below cut-off group (110) | 70 (63.6) | 40 (36.4) | 0.474 | 0.58 | 0.13 | 2.56 |
| Above cut-off (11) | 8 (72.7) | 3 (27.3) |  |  |  |  |

\*Safe water: filtered/boiled/ deep tube-well water; \*\*Unsafe drinking water: Tap water/ municipal supply/ pond water; <sup>§</sup>Per capita income USD 1751/- to date

Mean (±SD) age (years) of HEV IgM positive women were 26.03±6.8, whereas it was 29.09±9.9 for the negative cases. When HEV IgM positive patients were further divided based on pregnancy status, a comparable distribution (pregnant, n=37 and non-pregnant women, n=41) was found. The average duration of jaundice for HEV IgM positive cases was 13.68±9.27 days and it was significantly lower than that of the negative cases (21.12±16.03 days) (p=0.008), which further supports the acute status of the patients (Table 1b). Regarding biochemical parameters, serum ALT level, a potential liver damage marker was significantly higher in the HEV IgM positive cases than in the negative cases (p<0.0001). Similarly, the serum bilirubin and AST levels were also significantly higher in the positive group than the negative group, with the p-values of 0.027 and <0.0001, respectively (Table 2).

**Table 2.**
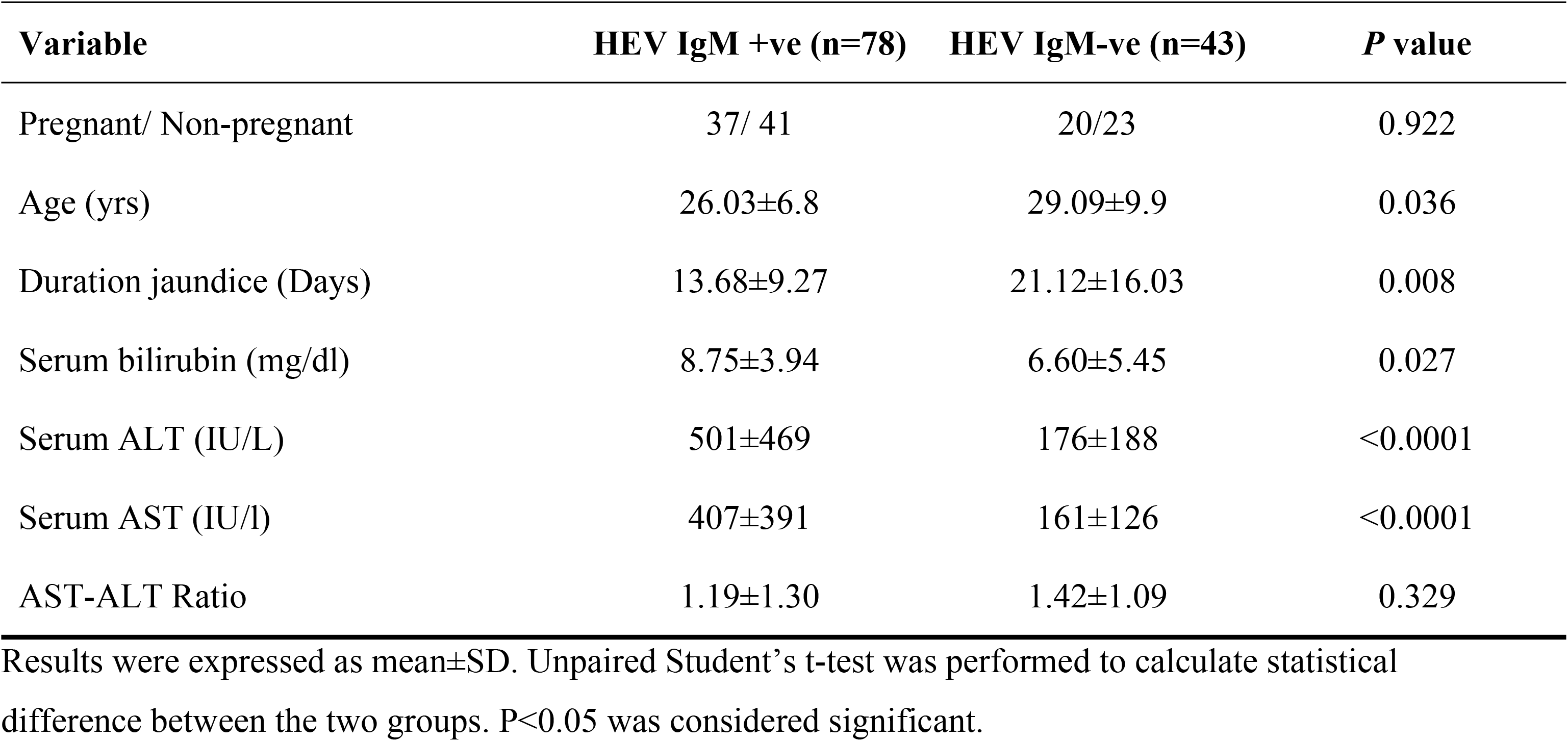
Pregnancy status, age, duration of jaundice and biochemical variables of the study subjects in respect to HEV IgM antibody status.

The HEV IgM positive cases were further categorized into 2 groups: early acute (duration of jaundice ≤14 days) and late acute phases (duration >14 days) of hepatitis [12]. Accordingly, among the 78 HEV IgM positive acute hepatitis cases, 48 (20 pregnant and 28 non-pregnant) were in early acute and 30 (17 pregnant and 13 non-pregnant) were in late acute phase of disease. Next, we wanted to compare whether there were any differences in ALT and AST levels, log HEV-RNA copy numbers and log HEV IgG sample index (SI) values between early and late acute phases of jaundice. As expected, serum ALT, AST and Log HEV viral copy number were significantly higher in early acute phase patients than in the late acute phase patients (p<0.0001, p=0.0036 and p<0.0001, respectively) (Fig 1A, 1B, 1C). Conversely, serum log HEV IgG SI value was significantly higher in late acute phase compared to early acute phase of jaundice (p<0.0001) (Fig 1D). The findings therefore suggest that an increase in IgG antibody levels at the late acute phase of HEV-induced jaundice might contribute to reduced hepatocellular damage resulting in production of lower levels of ALT, AST and log viral copy number, and thus demonstrating the protective role of IgG during HEV infection.

**Fig 1.**
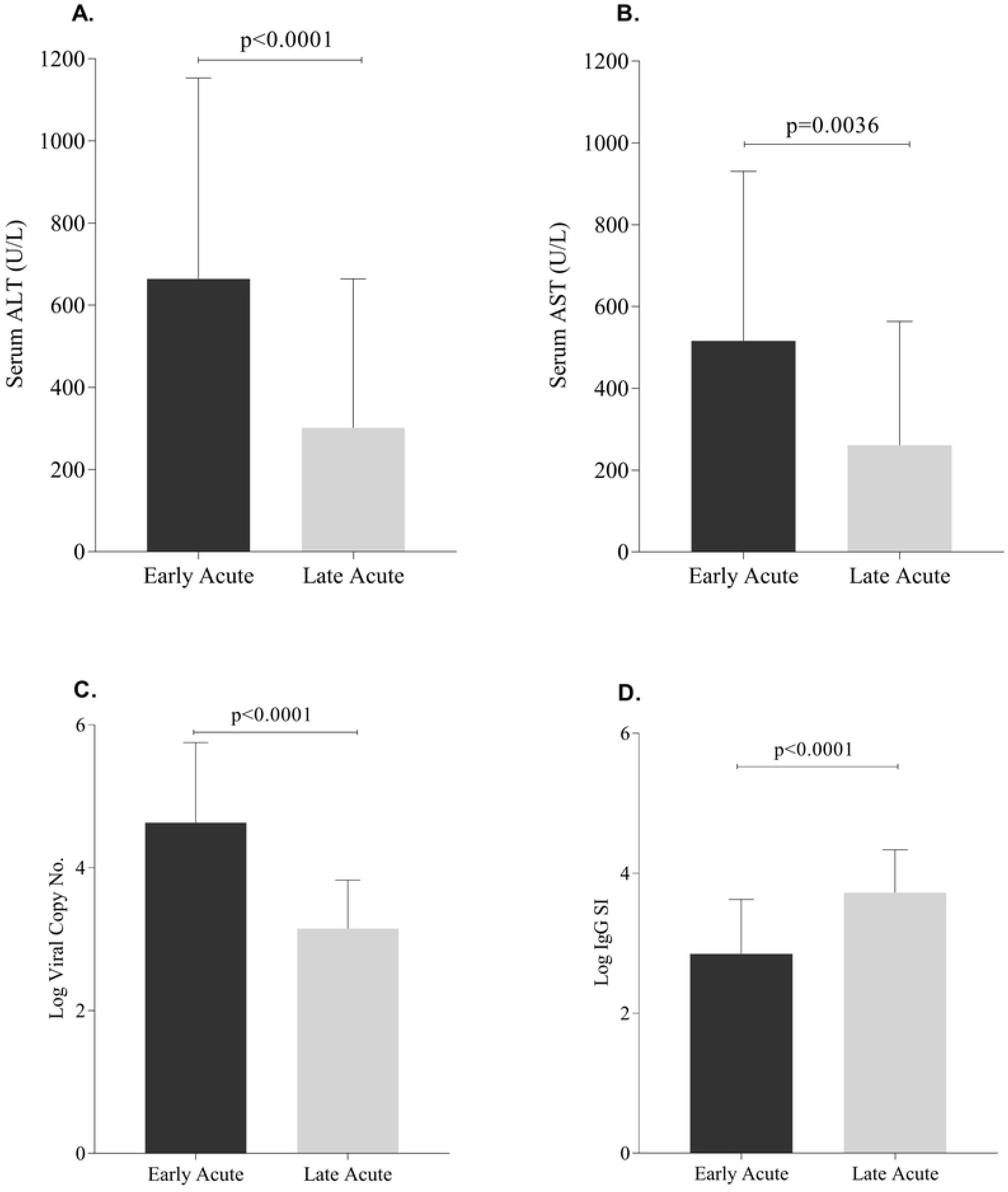
Comparison of (mean±SD) (A) serum ALT, (B) serum AST, and (C) log HEV copy number and (D) log HEV IgG SI values between early acute and late acute phases of HEV jaundice due to HEV infections. A p value <0.05 was considered significant.

### Correlation analysis between log viral copy number and log HEV IgG SI

Irrespective of pregnancy status, HEV-RNA log viral copy numbers and log HEV IgG SI value were inversely correlated in early and late acute phases of hepatitis. Although these two parameters did not show any significant correlation in early acute stage in pregnant women (r= −0.2278, p=0.3793) (Fig 2A), there was a significant negative correlation in non-pregnant counterpart (r=−0.5065, p=0.0320) (Fig 2B). On the other hand, significant negative correlations were observed between log viral copy number and log IgG SI values in the late acute phases of jaundice for both pregnant (r= −0.7971, p=0.0002) (Fig 2C) and non-pregnant women (r= −0.9117, p=0.0002) (Fig 2D). The result, therefore, suggests that correlation between HEV copy number and HEV IgG SI value depends on the intensity of the IgG immune responses during early and late acute phases of jaundice.

**Fig 2.**
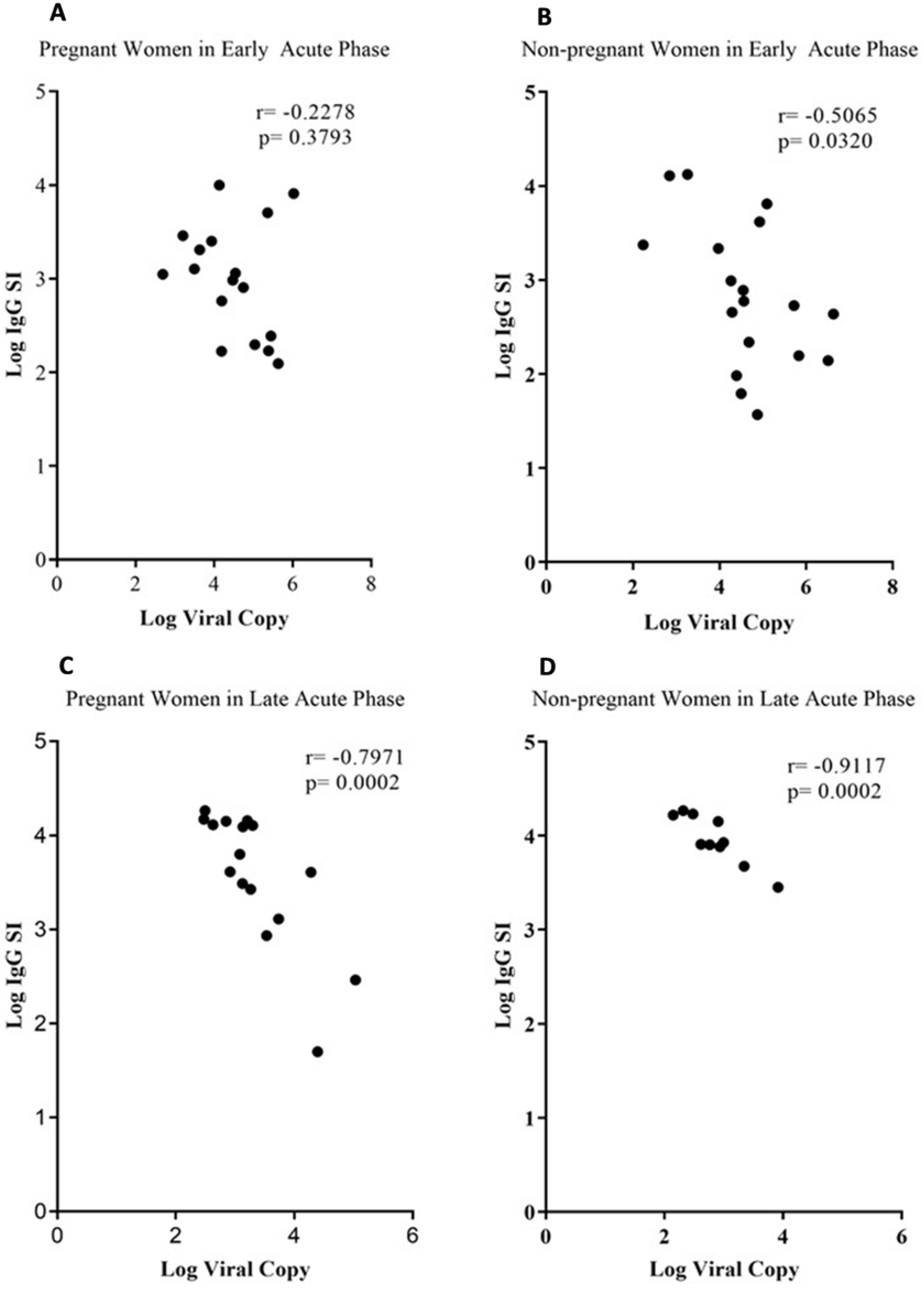
Pearson’s Correlation analysis between log HEV copy number and log HEV IgG SI value for pregnant and non-pregnant women in early and late acute phases of jaundice. (A) Log HEV copy number vs log HEV IgG SI of pregnant women in early acute phase; (B) Log HEV copy number vs log HEV IgG SI of non-pregnant women in early acute phase; (C) Log HEV copy number vs log HEV IgG SI of pregnant women in late acute phase; (D) Log HEV copy number vs log HEV IgG SI of non-pregnant women in late acute phase. A p value <0.05 was considered significant.

### Log viral copy and log IgG SI in pregnant vs non-pregnant

The HEV log viral copy number and log IgG SI value were compared between pregnant and non-pregnant women. As it is seen in Fig 3A and 3C, there were no differences in log viral copy number and log IgG SI value in early acute phase of jaundice between the pregnant and non-pregnant women (p=0.7905, p=0.6279, respectively). On the other hand, although log viral copy number in pregnant women was significantly higher than that of the non-pregnant counterpart (p=0.0196) (Fig 3B) in the late acute stage of jaundice, log HEV IgG SI value was significantly lower in the pregnant than in the non-pregnant women (p=0.0303) (Fig 3 D). The result, therefore, suggests that the increase in HEV copy number in the late acute phase of jaundice in pregnant women compared to the non-pregnant women might be due to a weak IgG immune response during pregnancy (Fig 3B, 3D).

**Fig 3.**
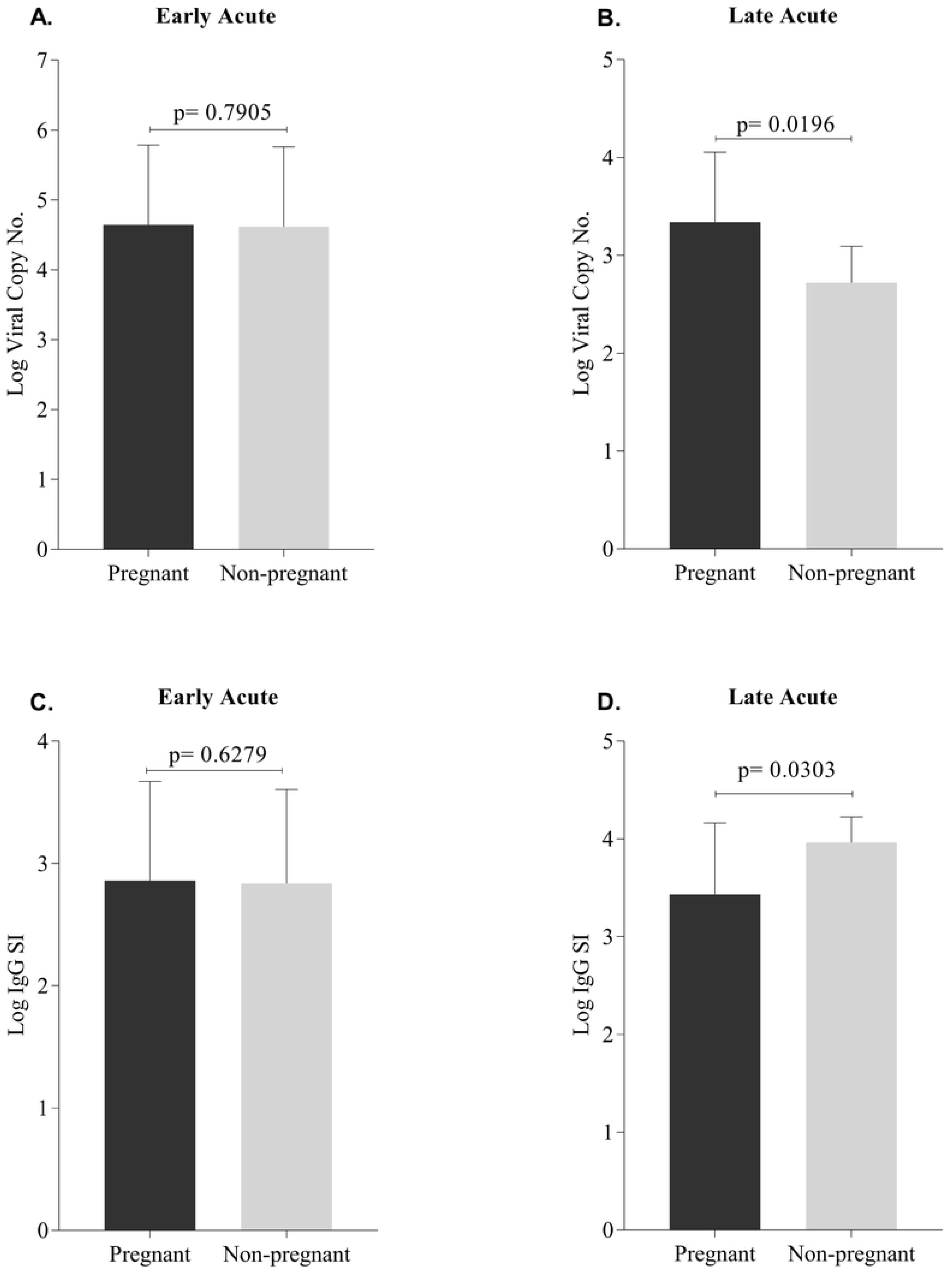
Comparison of Log HEV copy number and log HEV IgG SI value between pregnant and non-pregnant women with hepatitis in early (duration of jaundice ≤14 days) and late (duration of jaundice >14 days) phases of the disease. (A) Log HEV copy number for pregnant vs non-pregnant women in early acute phase; (B) Log HEV copy number for pregnant vs non-pregnant women in the late acute phase; (C) Log HEV IgG SI value for pregnant vs non-pregnant women in early acute phase; (D) Log HEV IgG SI value for pregnant vs non-pregnant women in the late acute phase. A p value <0.05 was considered significant.

The overall data suggest that HEV IgG has neutralizing effect on HEV-RNA and IgG immune response to HEV hepatitis in immunocompromised women during pregnancy.

### Anti-HEV IgG Avidity Test

We further examined if there were differences in anti-HEV IgG avidity between pregnant and non-pregnant women with HEV acute hepatitis. The anti-HEV IgG avidity test resulted in a significantly lower level of avidity index in the pregnant group compared to the non-pregnant patients (p=0.0017) (Fig 4). The findings indicate that more cross-reactive IgG antibodies are produced in pregnant women with acute hepatitis E than in the non-pregnant women with acute hepatitis E.

**Fig 4.**
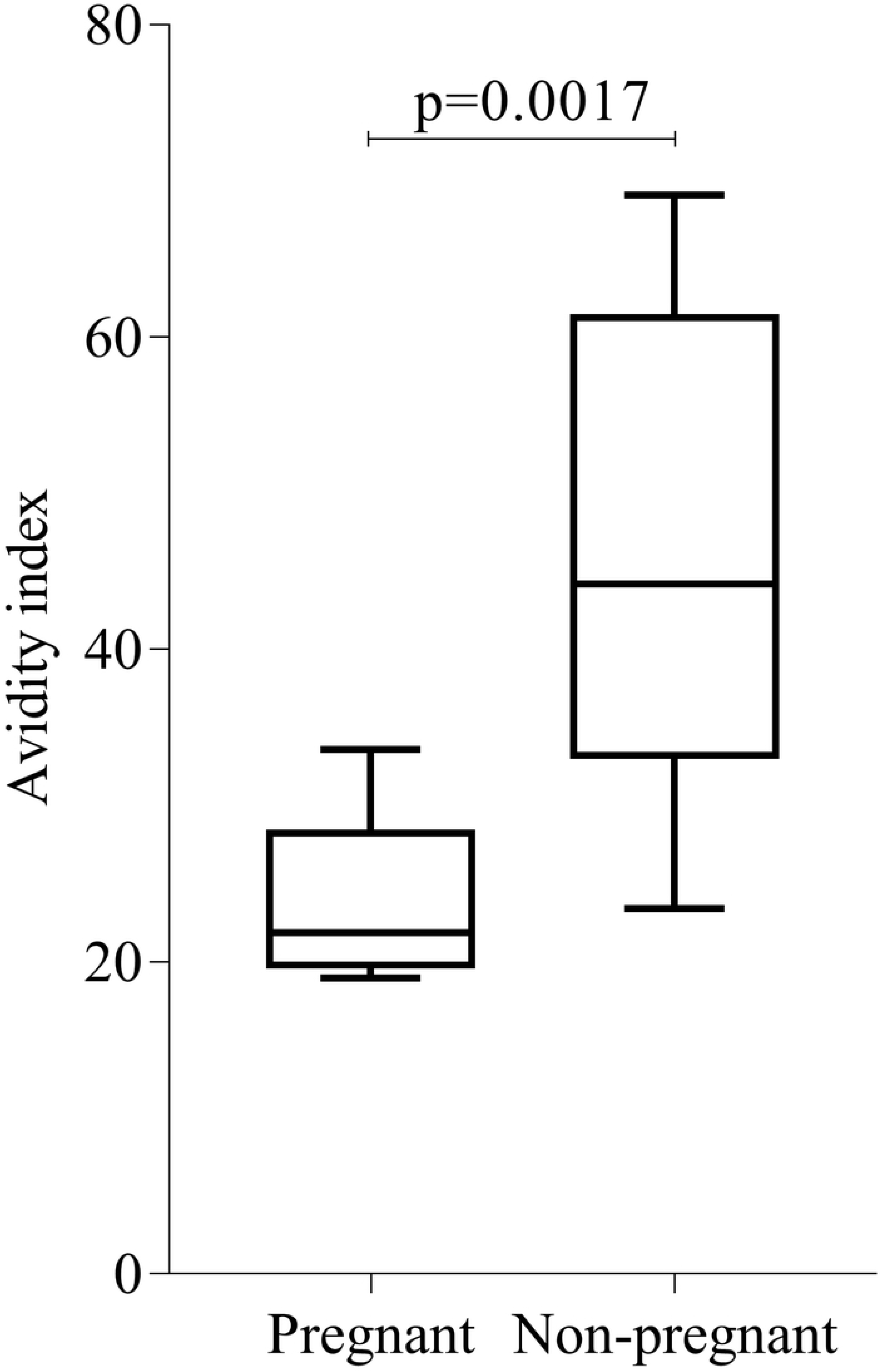
Comparison of IgG avidity index between pregnant and non-pregnant patient. A p value <0.05 was considered significant.

### Virial genotype determination

Sanger sequencing of HEV ORF2-specific PCR products and subsequent analysis by alignment with reference sequence showed that all the study samples that had been analyzed were HEV Genotype 1.

## Discussion

This is the first study demonstrating that virus-specific IgG immune response against acute HEV hepatitis are weaker in the pregnant women than in the non-pregnant women. Both animal and human studies demonstrated HEV IgG as protective against HEV infections [11–13] Since pregnant women had been reported to be vulnerable to HEV hepatitis, the present study explored whether HEV IgG immune responses during pregnancy were immunocompromised, which in turn, could contribute to vulnerability of pregnant women to acute HEV hepatitis.

All the commercially available HEV IgG ELISA kits are qualitative and as a result, it is not feasible to measure IgG immune responses against HEV virus by quantitative measurement of IgG using these commercially available IgG ELISA kits. However, to understand IgG immune responses without its quantitative measurement in HEV hepatitis, IgG antibody indices were measured to determine the fold-change of IgG HEV-infected sera by comparing its index with that of positive control, which was provided with the kit. The calculation of antibody indices could help to determine the role of HEV IgG in elimination of viruses and its association with reduced HEV load. The findings of significant inverse association between IgG antibody indices and HEV load in the late phase of jaundice in both pregnant and non-pregnant women could be considered as an evidence that IgG antibody, which was produced during HEV hepatitis, could render protection to HEV hepatitis. Furthermore, higher IgG indices, which were associated with lower HEV load, had significantly lower levels of liver injury markers including ALT, AST, and bilirubin, which indicates that IgG-mediated neutralization of HEV could improve the HEV-induced liver injury and jaundice. The HEV IgG levels in the present study was determined by using the kits from Beijing Wantai Biologicals, China, that was directed against recombinant HEV ORF2 protein antigen which was immunogenic and seemed to induce antibodies that possessed neutralizing effect on HEV and exert protective role.

Pregnant women have been reported to be more vulnerable to HEV infections, demonstrating 58% of maternal mortality in HEV-induced acute hepatitis in hospital care settings in Bangladesh [3]. So, we speculated that IgG production may be compromised in pregnant women with acute HEV hepatitis as a result of the disturbance of of Th2 cytokine bias during pregnancy [14] provided that IgG could possess protective function against HEV infections. To our expectation, pregnant women in their late phase of jaundice did have significantly lower levels of serum IgG against higher viral loads than their non-pregnant counterparts, which demonstrates weak IgG immune responses to HEV hepatitis during pregnancy. Compromised HEV IgG immune responses in pregnancy with HEV hepatitis was further supported by the presence of low level of IgG antibodies in the sera of early acute phase of jaundice of pregnant women and this phase of hepatitis was marked by higher levels of serum bilirubin, ALT, and AST as well as log of viral copy number, further demonstrating the compromised responses of humoral immunity due to lack of IgG antibodies during pregnancy with early acute HEV hepatitis. Compromised immune responses during pregnancy with acute hepatitis E is supported by a number of studies where they found generalized immune suppression with Th1/Th2 imbalance compared to the non-pregnant women with HEV [2, 14]. A state of maternal immune tolerance toward the fetus is characteristic in pregnancy [14]. Reduced T-cell activity with a concomitant reduction in cytokine production results in a dominating Th2 immune response in pregnancy that is interrupted by HEV infection, which strongly supports our finding that there was low level of IgG in late acute stage of HEV-induced jaundice [15].

Higher vulnerability to HEV infection during pregnancy may be associated with high levels of steroid hormones (estrogen, progesterone and human chorionic gonadotrophin), which are assumed to promote viral replication^2^. These steroid hormones have direct inhibitory effect on hepatic cells that may predispose to hepatic dysfunction or failure resulting in increased bilirubin, ALT, AST and high viral load upon exposure to infectious pathogens.

In the present study IgG antibody response was observed against recombinant HEV ORF2 antigen. Reports suggest that E2s domain (a.a455–a.a.602) within the ORF2 exhibit high level of immunogenicity and this peptide can be considered for manufacturing a candidate vaccine [16]. Because of the immunological potential and protective role of IgG antibodies against HEV, it can be passively transferred during outbreak of HEV with an aim to get neutralizing effect on HEV.

The study had a number of limitations. Although initially 153 patients (both pregnant and non-pregnant) with acute hepatitis were enrolled, we could not analyze all these samples because we needed exclusion of HAV, HBV, HCV positive and negative HEV (both IgM and IgG) cases. Finally, we could analyze only 121 samples with HEV acute hepatitis. Two pregnant patients in their third trimester succumbed to death and they both had antibodies (both IgM and IgG) to HEV. The possible reason of these deaths during third trimester may be due to interplay between host immunity and HEV. During pregnancy, high cytokine levels secreted by the HEV-specific antigen-stimulated peripheral blood mononuclear cells (PBMCs) coupled with Th1/Th2 cytokine imbalance could cause fulminant hepatic failure in pregnant women of later trimester [12]. However, we could not evaluate the cytokines levels (especially IFN-γ and IL-4 etc.) because of the unavailability of the reagents and this was another limitation of this study.

HEV IgG measurement in the present study was not an exact quantification, rather it was an alternative approach to obtain information of fold increase of HEV IgG in respect of positive control (when positive control absorbance was between 1.7 - 2.0) [10] that could help us to observe the correlation of true IgG levels with HEV viral load and other liver damage markers like serum ALT and AST.

## Abbreviations

CI: Confidence Interval
ELISA: Enzyme Linked Immunosorbent Assay
IFN-γ: Interferon-gama
IL-4: Interleukine-4
ORF2: Open Reading Frame 2
RT-PCR: Reverse Transcription-Polymerase Chain Reaction
SI: Sample Index
Th1: T helper 1
Th2: T helper 2

